# ASAP-ID: Proximity labelling with small tags

**DOI:** 10.1101/2025.08.20.671413

**Authors:** Ruohua Lyu, Kiersten M. Ruff, Catherine S. Palmer, Alejandra E. Ramirez, Angelique R. Ormsby, Daniel J. Scott, Rohit V. Pappu, Diana Stojanovski, David A. Stroud, Danny M. Hatters

**Affiliations:** Department of Biochemistry and Pharmacology and Bio21 Molecular Science and Biotechnology Institute, 30 Flemington Road. The University of Melbourne, Victoria 3010 Australia; Department of Biomedical Engineering, Center for Biomolecular Condensates, Washington University in St. Louis, St. Louis, MO 63130, USA; Materials Characterisation & Fabrication Platform. The University of Melbourne, Parkville, VIC 3052, Australia; Florey Institute of Neuroscience and Mental Health, The University of Melbourne, Parkville, VIC 3052, Australia; Murdoch Children’s Research Institute, Melbourne, VIC 3052, Australia; Victorian Clinical Genetics Services, Royal Children’s Hospital, Melbourne, VIC, 3052, Australia

## Abstract

Biotinylation-based proximity labelling methods are valuable for discovering protein-protein interactions within cellular systems. However, one limitation of these approaches is that most require fusing the target protein with the enzyme that biotinylates nearby proteins (i.e., TurboID or APEX2), which risks sterically disrupting the protein’s native function. Here, we present a method designed to reduce the steric impact of these fusions and offer greater flexibility in labelling modalities. The method, Antibody and Small-tag Assembly on Proteins for Interaction Detection (ASAP-ID), involves a bipartite system. Target proteins are fused to a small peptide antigen that recruits TurboID or APEX2 fused to an antibody directed to the antigen. Using two different antigen/antibody systems (SunTag and MoonTag), we show that ASAP-ID can specifically label human Lamin A in cells. The method works when the target protein and nanobody are co-expressed together *in cis* (ASAP-ID^IC^). We also demonstrate that the approach works when the antibody fusion is added *in trans* to fixed cells post-expression (ASAP-ID^IT^). ASAP-ID^IT^ identified more than 448 known and previously undescribed interactors of lamin. We further used ASAP-ID^IT^ to study how ALS-mutant profilin 1 affected its interactome. The method identified proteins involved in protein quality control that correlated with aggregation propensity. Moreover, the different mutants showed variation in the cellular location where aggregates formed. ASAP-ID^IT^ revealed preferences for mitochondrial proteins for the two profilin mutants that tend to aggregate in the cytoplasm, C71G and M114T, and nuclear proteins for a mutant more prone to nuclear aggregation. These findings position ASAP-ID as a powerful addition to the proximity labelling toolkit, capable of probing subtle differences in interactomes in a less invasive manner.

## Introduction

Defining the interacting partners of proteins in cells and tissues is one of the essential goals in biomolecular research. A leading approach to doing this involves proximity labelling, which involves closely localised or transiently interacting proteins being biotinylated for subsequent capture and detection. Most of the proximity labelling methods require the fusion of a catalytically promiscuous biotin ligase or peroxidase to the protein of interest (1). When substrate is added, biotinylation of proteins occurs within several nanometres of proximity in live cells, which can be captured by streptavidin affinity for identification by mass spectrometry (2). One of the first-developed proximity ligation approaches, BioID, used a variant of *Escherichia coli* biotin ligase BirA* (3). BioID has a relatively slow catalytic activity, often requiring hours in the presence of substrate for labelling to be sufficient for identification of interactors. However, fast-acting variants of BirA*, for example TurboID, have improved the reaction time to as low as 10 minutes (2). An alternative approach for faster labelling involves APEX2, an engineered soybean ascorbic acid peroxidase derivative that can label proteins within 30 seconds in the presence of hydrogen peroxide and substrate (4).

Both BirA* and APEX2 derivatives are fused to target proteins, which raises the potential for steric effects that hinder the target protein’s function and interactions. BirA* variants used in BioID and TurboID are about 38 kDa (2, 3). Efforts to reduce their size have brought them down to about 20 kDa, but this comes at the expense of decreased activity or protein stability (5). APEX2 is smaller than BirA* but still relatively large at 27 kDa (6).

One notable challenge we aimed to address was the requirement for an enzyme to be directly fused to a protein of interest. Here, we describe a proximity labelling strategy that utilises an epitope-antibody system to recruit enzymes to a target protein fused to a small (15-19 amino acid) epitope tag. We validate this approach using Lamin as a well-characterised benchmark for proximity labelling, and then apply it to determine how disease-relevant mutations in profilin 1, which lead to protein aggregation, alter the set of proteins in proximity to profilin. Of particular interest was the method’s ability to perform proximity labelling of target proteins on fixed cells by adding the proximity labelling enzyme *in trans*, offering unique methodological advantages for research applications.

## Experimental Procedures

### Cell culture

The human embryonic kidney (HEK) 293T and HeLa cell lines were obtained from the American Type Culture Collection (ATCC, Manassas, Virginia). HEK293T cells were cultured in Dulbecco’s Modified Eagle’s Medium (DMEM) supplemented with 2 mM L-glutamine and 10% v/v fetal bovine serum (complete DMEM). HeLa cells were cultured in DMEM supplemented with 10% v/v fetal bovine serum and 1% v/v penicillin-streptomycin (Thermo Fisher Scientific). To detach cells for passage, 0.05% w/v Trypsin and 0.02% w/v EDTA in PBS were used. Cells were maintained in a humidified incubator at 37 °C with 5% v/v atmospheric CO_2_.

### Plasmids and Cloning

The plasmid sequences developed for the project are listed in **Table S1**. Human lamin A, APEX2, MoonTag, MoonTag nanobody, wild type and mutant profilin1 cDNA were synthesised commercially (Twist Bioscience). The cDNA of the anti-SunTag scFv, HA tag, and GB1 were amplified by PCR from the plasmid pHR-scFv-GCN4-HaloTag-GB1-NLS-dWPRE (Addgene, #106303 (7)). The cDNA of TurboID was amplified from pcDNA V5-TurboID-NES (Addgene, #107169 (2)) by PCR. The scFv-HA-TurboID/APEX2-GB1 plasmids were generated from the pHR vector using restriction enzyme-mediated cloning. SunTag cDNA was amplified from pcDNA4TO-5xGCN4_v4-kif18b-24xPP7 by PCR (Addgene, #74927 (8)). The mCherry-SunTag-lamin and mCherry-Lamin expression plasmids were created by replacing GFP with mCherry in the pEGFP-C2 plasmid (Clontech Laboratories), and inserting the SunTag and lamin cDNA using restriction enzyme cloning. HiFi DNA assembly (New England Biolabs) was used to replace SunTag with MoonTag. The mCherry was swapped with APEX2 to generate the APEX2-lamin plasmid via restriction enzyme cloning. For the SunTag-tagged profilin1 (PFN1) expression plasmids, EGFP was replaced with SunTag or SunTag-HA sequences in the Gateway cloning destination vector (Addgene, #122844 (9)) using HiFi DNA assembly. The wildtype and mutant PFN1 sequences were commercially synthesised in a Gateway Entry vector (Twist Bioscience). PFN1-SunTag and PFN1-SunTag-HA were transferred into Gateway expression vectors using the Gateway LR Clonase Enzyme mix (Thermo Fisher Scientific).

### Transfection

Cells were seeded either 4.5 × 10^4^ cells into 8-well µ-slides (Ibidi) for immunofluorescence studies, 5 × 10^5^ cells into 6-well plates (Corning) for Western Blot, 1.1 × 10^6^ cells into each T25 flask (Thermo Fisher Scientific) for proteomic experiments or 4 × 10^6^ cells into T75 flasks (Thermo Fisher Scientific) for antibody purification. Cells were cultured in complete DMEM for 18 hours before transfection. Cells were transfected with plasmids using Lipofectamine 3000 (Thermo Fisher Scientific), following the manufacturer’s guidelines. Five hours after transfection, the media was replaced with complete DMEM.

### Methanol Fixation

Twenty-four hours after transfection, cells were washed with phosphate-buffered saline (PBS) and then fixed with methanol (pre-chilled at −20 °C overnight) at –20 °C for 10 minutes. The methanol was removed, and the cells were rinsed three times with PBS.

### Paraformaldehyde Fixation

Twenty-four hours after transfection, cells were rinsed with PBS and then fixed with 4% w/v paraformaldehyde at room temperature for 10 minutes. The paraformaldehyde was removed and the cells were incubated with 50 mM ammonium chloride at room temperature for 10 minutes and then washed three times with PBS. For immunofluorescence, the cells were permeabilised with 0.1% v/v Triton X-100/PBS at room temperature for 10 minutes and washed three times with PBS.

### ScFv/Nanobody-APEX2 Purification

Cells transfected with scFv-HA-APEX2-GB1 or Nanobody-HA-APEX2-GB1 plasmids were lysed using a Triton X-100-based lysis buffer (20 mM Tris-HCl, 150 mM NaCl, 10% v/v Glycerol, 2 mM EDTA, 1% v/v Triton X-100, 1 mM phenylmethylsulfonyl fluoride, and protease inhibitor cocktail (Sigma, #11836170001), pH 8.0) on ice for 10 minutes. Cell lysates were extruded through a 30 G syringe ten times, followed by centrifugation at 21,000 g for 10 minutes at 4 °C. The supernatant was incubated with pre-washed anti-HA magnetic beads (Thermofisher) at room temperature for 3 hours while rotating. The prewash involved rinsing the beads three times with TBS-T (Tris-buffered saline (TBS) 20 mM Tris, 150 mM NaCl containing 0.05% v/v Tween-20, pH 8.0). The anti-HA magnetic beads were washed three times with TBS-T, and the APEX2-antibody was eluted using 2 mg/mL HA peptide (in TBS, MedChemExpress) by incubating at 37 °C for 20 minutes while rotating. The supernatant was snap-frozen in liquid nitrogen and stored at –80 °C.

### In-trans APEX2 Biotinylation

The purified scFv-APEX2 or nanobody-APEX2 was applied to fixed cells and incubated overnight at 4 °C with shaking. The cells were rinsed three times with PBS and then incubated in complete DMEM containing 100 µM biotin-phenol (Sigma) for 10 minutes at room temperature. The biotinylation reaction was initiated by adding H_2_O_2_ to a final concentration of 1 mM for 1 minute, followed by quenching with an *in-trans* quenching buffer (10 mM sodium ascorbate, 5 mM Trolox in PBS). Cells were subsequently washed twice with the *in-trans* quenching buffer.

### In-cis APEX2 Biotinylation

Twenty-four hours post-transfection, cells were rinsed once with PBS and incubated in complete DMEM containing 500 µM biotin-phenol (Sigma) for 30 minutes in a humidified incubator at 37 °C with 5% v/v atmospheric CO_2_. The biotinylation reaction was initiated by adding H_2_O_2_ to a final concentration of 1 mM for 1 minute and quenched by replacing the media with an *in-cis* quenching buffer (10 mM sodium azide, 10 mM sodium ascorbate, 5 mM Trolox in PBS). Cells were then washed with *in-cis* quenching buffer twice.

### In-cis TurboID Biotinylation

Twenty-four hours after transfection, cells were rinsed once with PBS and incubated in complete DMEM containing 500 µM biotin (Sigma) for 2 hours in a humidified incubator at 37 °C with 5% v/v atmospheric CO2. Cells were then washed with cold PBS (chilled on ice for 1 hour) three times.

### Immunofluorescence staining and imaging

For the biotinylation experiments, cells were blocked with Triton blocking buffer (0.1% v/v Triton X-100 and 5% w/v bovine serum albumin in PBS) for 1 hour at room temperature. Streptavidin-conjugated Alexa Fluor 488 (Thermo Fisher Scientific, #S11223) was added for 1 hour at room temperature (1:500 dilution in the Triton blocking solution). After two washes with PBS, cells were stained with Hoechst 33342 (1:1000 dilution in PBS, Thermo Fisher Scientific, #H3570) for 10 minutes at room temperature, followed by three more PBS washes. Immunofluorescence images were acquired on a LSM880 confocal microscope (Zeiss) with a 60× objective.

For experiments not involving biotinylation, cells were blocked with blocking buffer (3% w/v bovine serum albumin in PBS) for 1 hour at room temperature and incubated with anti-SDHA antibody (1:500 dilution in the blocking buffer, Abcam, #AB14715) at room temperature for 1 hour. After two washes with PBS, cells were probed with an anti-HA antibody (1:500 dilution in blocking buffer, Cell Signalling, #3724S). After another two washes with PBS, cells were stained with Anti-rabbit Alexa Fluor 568 (1:1000 dilution in PBS, Thermo Fisher Scientific, #A-11011) and Anti-mouse Alexa Fluor 488 (1:1000 dilution in PBS, Thermo Fisher Scientific, #A-11001) respectively at room temperature for 1 hour, with two washes of PBS after each incubation. Immunofluorescence images were acquired on a Zeiss Elyra LSM880 confocal microscope (Zeiss) with a 63×/1.4 NA Plan-Apochromat oil immersion objective. Super-resolution structured illumination microscopy (SIM) was performed on a Zeiss Elyra 7 Lattice SIM (Zeiss) with a 63×/1.4 NA Plan-Apochromat oil immersion objective and a pco.edge sCMOS 4.2 CL HS camera. SIM reconstructions were performed with Zeiss ZEN Black 3.1 SR software with the Lattice SIM2 processing function.

### Enrichment with streptavidin and sample preparation for mass-spectrometry

HEK293T cells were plated on T25 flasks and transfected with the lamin or PFN1 constructs. Cells were then processed as described above for the methanol or paraformaldehyde fixation and biotinylation steps. For methanol fixation, cells were resuspended in 0.5 mL of buffer containing 8 M urea, 2% w/v SDS, 0.1 M DTT and incubated for 10 minutes at room temperature, followed by adding 1 µL Universal Nuclease (Thermo Fisher Scientific) and another 10 minutes incubation at room temperature. The lysate was transferred into LoBind tubes (Eppendorf) and centrifuged at 21,000 g for 10 minutes at room temperature. The supernatant was transferred into new LoBind tubes and underwent the PD-10 desalting step. For the paraformaldehyde fixation, cells were lysed in 0.5 mL of the buffer (10 mM Tris-HCl, 140 mM NaCl, 1 mM EDTA, 0.5% w/v sodium deoxycholate, 1% v/v Triton X-100, 2% w/v SDS, 1 mM phenylmethylsulfonyl fluoride, and a protease inhibitor cocktail (Sigma), pH 8.0) for 10 minutes on ice. The lysate was transferred into LoBind tubes and boiled at 100 °C for 20 minutes, followed by incubation at 60°C for 2 h. The lysate was then centrifuged at 21,000 g for 10 minutes at room temperature. The supernatant was transferred into new LoBind tubes and desalted with PD-10 columns (Cytiva). Protein concentrations were determined by optical density (AU280 nm) on a Nanodrop (Thermo Fisher Scientific) using Bovine Serum Albumin for the standard curve. Magnetic streptavidin beads (Thermo Fisher Scientific) were washed with PBS twice. 600 µg of protein was incubated overnight with 100 µL of pre-washed magnetic streptavidin beads (Thermo Fisher Scientific) at 4 °C with rotation. The unbound protein fraction was collected. Magnetic beads were washed twice with 1 mL of urea buffer (6 M urea, 100 mM Tris-HCl, pH 8.0) and once with 1 mL of PBS containing 0.5% w/v SDS. The beads were then incubated with 200 µL of PBS containing 0.5% w/v SDS and 100 mM DTT at room temperature with shaking at 700 rpm. The beads were washed twice with 1 mL of urea buffer and incubated with 200 µL of urea buffer containing 50 mM chloroacetamide for 2 h at room temperature in the dark. Beads were washed once with 1 mL of urea buffer, once with 1 mL of PBS, and once with 1 mL of 50 mM ammonium bicarbonate (ABC), followed by a wash with 0.5 mL of 50 mM ABC. After washing, each sample was incubated with 100 µL of 50 mM ABC containing 1 µg of trypsin (Thermo Fisher Scientific) at 37 °C overnight. The digested fractions were collected and acidified to 1% v/v trifluoroacetic acid (TFA) by 10% v/v TFA. Stage tips were pre-packed with two layers of Empore SPE Disks (Merck, #66884-U) membrane and activated with 50 µL of 100% v/v acetonitrile (ACN) by centrifuging at 1,500 g, 22 °C until all liquid went through. Tips were washed with 50 µL of 0.1% v/v TFA, 2% v/v ACN. Samples were loaded onto the tips and centrifuged as above. Next, tips were washed with 100 µL of 0.1% v/v TFA and 2% v/v ACN again and transferred into new LoBind tubes into which peptides were eluted with 100 µL of 0.1% v/v TFA, 80% v/v ACN, dried by SpeedVac, and stored at –80°C.

### Mass spectrometry data acquisition

LC-MS/MS was performed on an Orbitrap Ascend mass spectrometer (Thermo Scientific) equipped with a nanoflow reversed-phase-HPLC (Ultimate 3000 RSLC, Dionex) fitted with an Acclaim Pepmap nano-trap column (Dionex—C18, 100 Å, 75 µm× 2 cm) and an Acclaim Pepmap RSLC analytical column (Dionex—C18, 100 Å, 75 µm× 50 cm). The tryptic peptides were injected into the enrichment column at an isocratic flow of 5 µL/min of 2% v/v ACN containing 0.1% v/v formic acid for 5 minutes, applied before the enrichment column was switched in line with the analytical column. The eluents were 5% DMSO in 0.1% v/v formic acid (solvent A) and 5% DMSO in 100% v/v ACN and 0.1% v/v formic acid (solvent B). The flow gradient was (i) 0-6min at 3% B, (ii) 6-7min, 3-4% (ii) 7-82 min, 4-25% B (iii) 82-86min 25-40% B (iv) 86-87min, 40-80% B (v) 87-90min, 80-80% B (vi) 90-91min, 80-3% and equilibrated at 3% B for 10 minutes before the next sample injection. The mass spectrometer was operated in data-dependent acquisition mode, whereby complete MS1 spectra were acquired in a positive mode at 120,000 resolution. The ‘top speed’ acquisition mode (3 s cycle time) on the most intense precursor ion was used (i.e., charge states of 2 to 7). MS/MS analyses were performed by 1.6 *m/z* isolation with the quadrupole, fragmented by HCD with a normalised collision energy of 30%. MS2 fragmented ion spectra were acquired at 15,000 resolution. Dynamic exclusion was activated for 30 s, and the AGC target was set to standard with auto maximum injection mode.

### Mass spectrometry data analysis

The raw files were searched using MaxQuant (v. 2.4.3.0) with the UniProt human proteome as the reference (downloaded in January 2023). The enzyme specificity was set as trypsin, and the maximum number of missed cleavage sites permitted was two. Variable modifications were used for all experiments: oxidation (M), acetylation (Protein N-term). A fixed modification used for all experiments was carbamidomethyl (C). Label-free quantification was applied without normalisation. 1% FDR was used. The match between runs setting was selected in the identification parameters. The mass tolerance for precursor ions was 20 ppm, and the mass tolerance for fragment ions 20 ppm.

### Western blot

The input and the unbound protein fractions collected during the enrichment using magnetic streptavidin beads (Thermo Fisher Scientific) as described above were analysed by Western blot. Samples were mixed with SDS loading dye and dithiothreitol (DTT). The samples were boiled at 95°C for 10 minutes before electrophoresis. The boiled samples were loaded onto a 12% fast-cast stain-free SDS-PAGE (Bio-Rad) and run at 130 V for 1 hour. After electrophoresis, proteins were transferred to a 0.2 µm pore size PVDF membrane using the iBlot transfer system (Thermo Fisher Scientific) for 7 minutes at 20 V. The membranes were blocked with a Western Blot blocking buffer (3% w/v BSA in PBS) at room temperature for 1 hour, then incubated with streptavidin-HRP (1:1000 dilution in the Western Blot blocking buffer, Thermo Fisher Scientific, #S911) at room temperature for 30 minutes. Following incubation, the membranes were washed in PBS-T (PBS with 0.05% v/v Tween-20) with shaking at room temperature for 30 minutes. The enhanced chemiluminescence (Bio-Rad) reagent was prepared according to the manufacturer’s instructions and applied to the membrane for protein detection. Images were captured using the ChemiDoc imaging system (Bio-Rad).

### Experimental Design and Statistical Rationale

For ASAP-ID and traditional proximity labelling constructs of APEX2-lamin fusion experiments, controls were established differently. In the traditional fusion approach, free APEX2 was expressed in cells as the background control and was compared to the APEX2-tagged protein of interest. In ASAP-ID, we compared the untagged protein against the epitope-tagged protein of interest. There were four biological replicates for each sample.

For the analysis of lamin proteomics data, the proteinGroups text file generated by MaxQuant was loaded into Perseus (v 2.0.10.0), with the LFQ intensity of each sample as the main variable. LFQ intensities were log2 transformed and filtered to remove rows where LFQ intensities were not reported in at least three SunTag-containing biological replicates in ASAP-ID experiments or at least three APEX2-tagged protein of interest biological replicates in the direct fusion experiments. The data was then normalised to the pyruvate carboxylase (PC) protein by subtracting the PC LFQ intensities row from all other LFQ intensities in the column. A default imputation was performed on non-SunTag-tagged samples in the ASAP-ID experiments and free APEX2 samples in the direct fusion experiments. For ASAP-ID, a two-tailed two-sample t-test was conducted on log2 LFQ intensities comparing the SunTag-tagged and non-tagged protein of interest. The significance threshold for the methanol ASAP-ID was set as FDR < 0.05 and S0 = 1.7; for the paraformaldehyde ASAP-ID, it was FDR < 0.05 and S0 = 4.8. This threshold was set to exclude all proteins detected in the non-tagged lamin samples on the assumption they represent background enrichment. For the traditional proximity APEX-lamin fusion method, a common threshold was applied: p-value < 0.05 and fold change > 2. Gene ontology terms were analysed using PANTHER GO-slim (v19.0) or STRING (v12.0) with default settings.

For the ASAP-ID on PFN1, the non-tagged wildtype and mutant PFN1 served as controls and were compared to the SunTag-tagged wildtype and mutant PFN1. Each sample had three biological replicates.

For the analysis of PFN1 proteomics data, the proteinGroups text file generated by MaxQuant was loaded into Perseus (v2.0.10.0), with the LFQ intensity of each sample as the primary variable. Log2 transformed LFQ intensities were filtered to exclude those not identified in at least two SunTag-containing biological replicates. The data was normalised to the PC protein by subtracting the PC protein intensity row from all rows. A default imputation was performed on non-SunTag-tagged samples. LFQ intensities were then smoothed using PERCEPT (10), a tool that enhances focus on statistically meaningful changes in the data. The scaled LFQ intensity log2 fold change after PERCEPT was loaded back into Perseus, followed by k-means clustering analysis. The optimal number of clusters was determined by calculating the inertia with the sklearn machine learning library in Python. Actual clusters were generated in Perseus using Euclidean distance. Gene ontology terms were analysed using PANTHER GO-slim (v19.0) or STRING (v12.0) with default settings.

For the analysis of immunofluorescence images of PFN1-expressing HeLa cells, PFN1 was quantified in cells expressing the C71G, M114T, and G118V mutants using ImageJ FIJI software. 25 to 38 cells from each variant were measured. The data were analysed statistically using GraphPad Prism (v10). For two-sample comparisons, the data were analysed by an unpaired Student’s t-test, and the data passed the Normality test. For multiple samples, we used one-way ANOVA with differences evaluated according to the two-stage linear step-up procedure of Benjamini, Krieger and Yekutieli (11), which was used because it is sensitive and well-suited to detect the true positives even when data are positively dependent. We used the C71G mutant as the reference sample for the test.

Surfaces of PFN1 variants and SDHA were generated from super-resolution images using Imaris software (v10.0). Imaris was also used to measure the distance between PFN1 and SDHA.

## Results

The logic of the Antibody and Small-tag Assembly Proximity-Interaction Detection (ASAP-ID) method is outlined in **Fig. 1A**. It involves fusing a small epitope tag (e.g., SunTag or MoonTag) to the target protein, which binds to an antibody derivative (e.g. nanobody or single-chain variable fragment (scFv) chain) fused to APEX2 or TurboID. The SunTag is a 19-amino-acid epitope for an scFv (12) and the MoonTag is a 15-amino-acid epitope for a camelid variable heavy domain of heavy chain nanobody (13). The antibody derivative fusion also contains the 6.2 kDa GB1 domain, derived from the immunoglobulin-binding domain of protein G (14), which is designed to suppress aggregation (12). A further element is a HA tag, which is used for purification of the antibody derivative fusion to APEX2 or TurboID by immunoaffinity chromatography. To apply ASAP-ID, cells are transfected to express the protein of interest fused to the small epitope tag. Cells can be co-transfected with the antibody-biotinylating enzyme fusion for *in-cis* labelling (ASAP-ID^IC^) (**Fig. 1B**).

**Figure 1.**
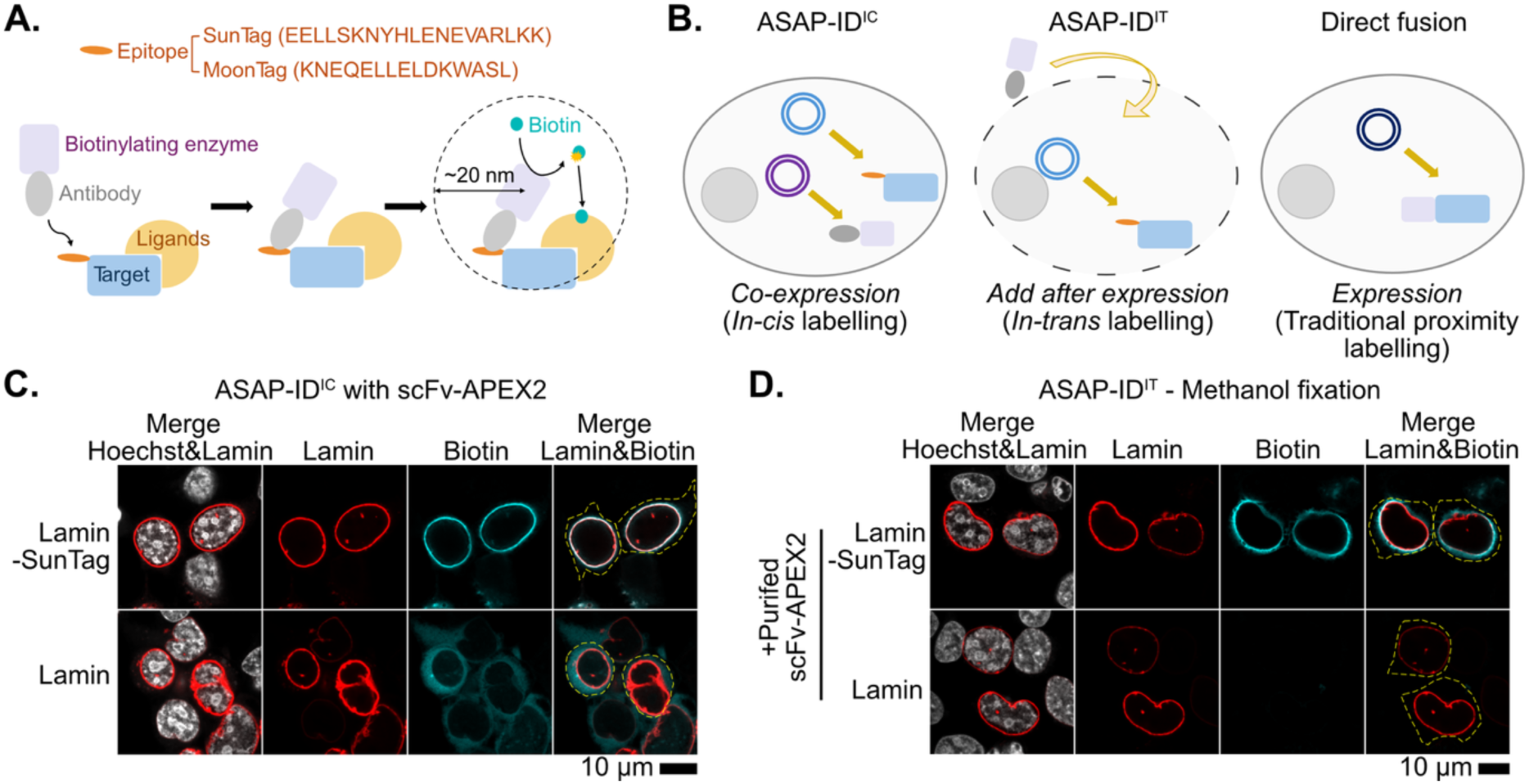
ASAP-ID can label proximal proteins of the target protein with biotin. **A.** Shown is a schematic of the ASAP-ID approach. The epitope can be either a SunTag or a MoonTag. The proximity ligase enzyme for biotinylating can be APEX2 or TurboID. The antibody, which is a nanobody or scFv, is fused to the biotinylating enzyme, allowing it to be recruited to the tagged bait protein and proximal molecules labelled with biotin. **B.** Illustration of *in-cis* or *in-trans* versions of ASAP-ID (ASAP-ID^IC^ and ASAP-ID^IT^ respectively) and the traditional fusion proximity labelling method. In ASAP-ID^IC^, the tagged bait protein and antibody-fused biotinylating enzyme are co-expressed in cells. In ASAP-ID^IT^, the tagged bait protein is first expressed in cells, and then the antibody-fused biotinylating enzyme is introduced to the fixed cells for subsequent labelling. In the traditional method, the bait protein is fused directly to the biotinylating enzyme. **C.** ASAP-ID^IC^ with SunTag and APEX2. The SunTag-tagged or non-tagged Lamin was co-transfected with scFv-APEX2 into HEK293T cells. Cells were then subjected to biotinylation and fixed with methanol. **D.** ASAP-ID^IT^ with methanol fixation. HEK293T cells were transfected with lamin constructs and then fixed with methanol, and the scFv-APEX2 fusion proteins were added *in-trans*. Antibody-APEX2 was either purified using HA antibody affinity chromatography or expressed in cells, where cell lysates were added.

Alternatively, cells can be transfected to express just the protein of interest-epitope tag fusion and fixed for *in-trans* labelling by application of the semi-purified antibody-APEX2 protein, a variation of the method we call ASAP-ID^IT^ (**Fig. 1B**). For ASAP-ID^IT^, the antibody-APEX2 can be added as crude cell lysate from cells transfected with antibody-APEX2 or as a partially purified product using HA-immunoaffinity chromatography. ASAP-ID contrasts to the traditional proximity labelling methods that involve fusing the biotinylating enzyme directly to the protein of interest (**Fig. 1B**).

To develop ASAP-ID, we first tested it on human lamin A (lamin), the target protein initially used to develop BioID (3). Lamin is a component of the nuclear lamina and is easy to visualise by immunofluorescence confocal microscopy in transfected cells, as it is localised almost exclusively at the nuclear membrane. When the scFv-APEX2 or scFv-TurboID was co-expressed with SunTag-tagged lamin (**Fig. 1C & Fig. S1A)**, specific labelling was evident at the nuclear membrane, indicating that the ASAP-ID^IC^ proximity labelling had worked. Without the SunTag, biotinylation appeared to be non-specific and was visible throughout cytosol.

Next, we evaluated whether the *in-trans* labelling was effective for ASAP-ID^IT^. HEK293T cells were transfected with SunTag-tagged lamin, fixed with either methanol or paraformaldehyde, and then treated, *in-trans*, with scFv-APEX2 semi-purified via HA-immunoaffinity chromatography or with crude cell lysate expressing scFv-APEX2. Both the semi-purified and unpurified scFv-APEX2 caused specific biotinylation of SunTag-tagged lamin compared to lamin lacking the SunTag (**Fig. 1D & Fig. S1B**). The semi-purified scFv-APEX2 in lysate maintained most of its activity after being stored as frozen aliquots for a month (**Fig. S2**). Specific labelling *in trans* was also observed with the MoonTag system (**Fig. S1C**), indicating that ASAP-ID^IT^ can easily be adapted to different epitope tagging systems. Moreover, sufficient semi-purified scFv-APEX2 can be obtained from ∼20 × 10^6^ cells and small-scale immunoprecipitation using anti-HA magnetic beads, which suggests this methodology will be widely and easily accessible to researchers.

Next, we assessed whether ASAP-ID^IT^ would work in a proteomics format. For these tests, we compared SunTag-lamin with untagged lamin. As a control, we performed a traditional proximity labelling experiment with lamin fused to APEX2 and compared it to APEX2 alone. As expected, ASAP-ID^IT^ resulted in a greater level of biotinylation when the SunTag epitope was present (**Fig. 2A & Fig. S3A**). Unexpectedly, labelling of APEX2 alone had a greater level of biotinylation than when directly fused to lamin (**Fig. S3B**). This indicates that free APEX2 in the cell exhibits a high level of non-specific reactivity to the proteome, but the *in-trans* labelling strategy of ASAP-ID^IT^ effectively avoids this because unbound APEX2 is washed away. In principle, this supports the idea that ASAP-ID^IT^ offers increased specificity for the target protein.

**Figure 2.**
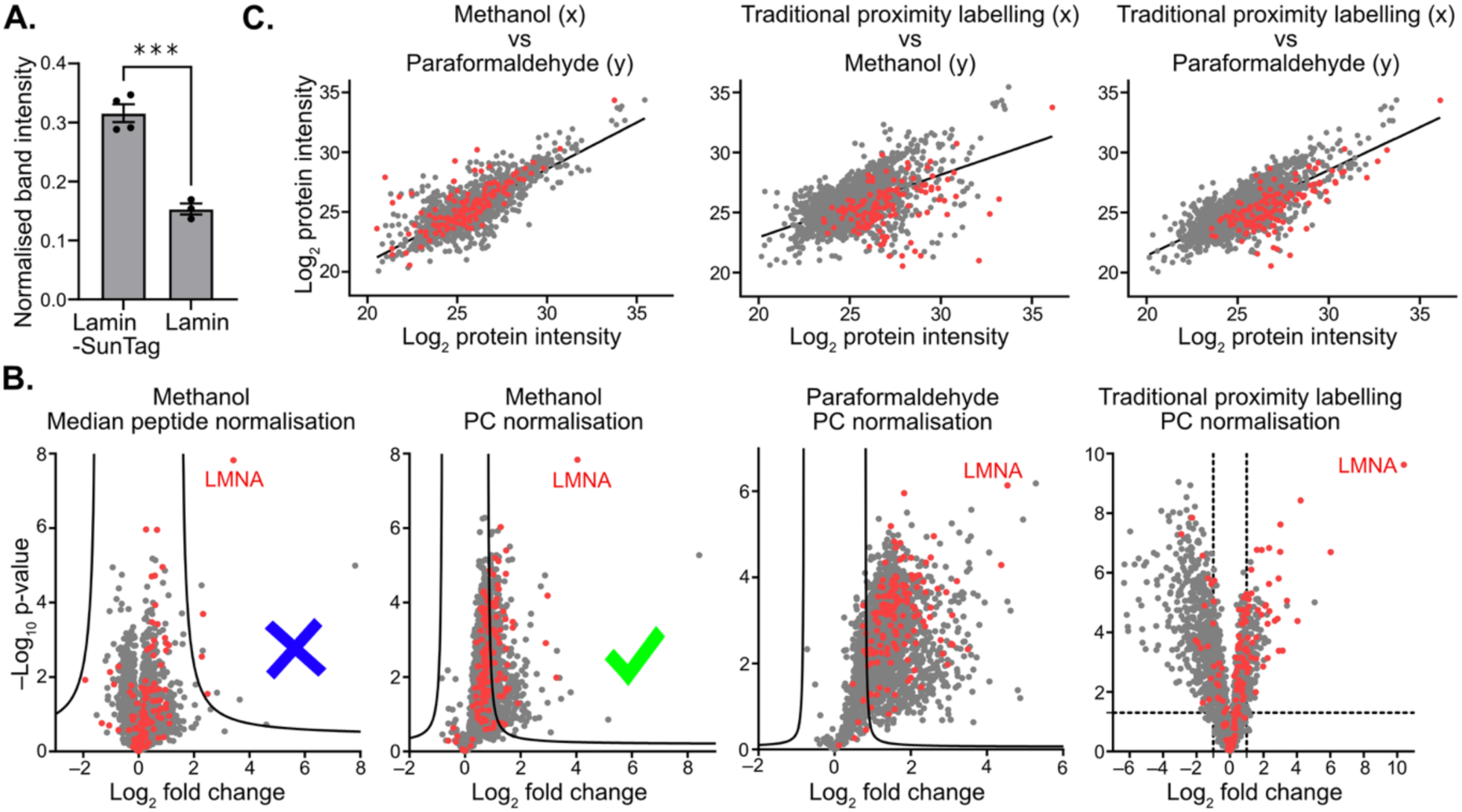
ASAP-ID identifies proximal proteins of lamin. **A.** Quantification of total biotinylated proteins on Western Blot of an ASAP-ID^IT^ conducted on HEK293 cells transfected with the shown constructs (blots in Fig S3A; data shows densitometry data; means and SEM. T-test result shown: ***, P-value = 0.0004). B. Volcano plots using different normalisation approaches. Labels above the graph depict the method of fixation for the ASAP-ID experiment, or the traditional proximity labelling method by expression of the APEX-lamin fusion (these cells are not fixed). Red dots represent previously established lamin interactors. **C.** Comparison of proteins identified by the different proximity labelling methods. The lines represent the linear regression models with R^2^ values of 0.6402 (methanol versus paraformaldehyde), 0.2902 (Traditional proximity labelling versus methanol), and 0.5739 (Traditional proximity labelling versus paraformaldehyde).

Quantitative analysis of biotinylation-based proximity labelling data requires careful consideration of the method used for abundance normalisation. In standard proteomics of cell lysates, protein abundances are typically normalised by median peptide counts, a method we call “median normalisation.” This approach presumes that total protein levels are similar across the datasets being compared, which is reasonable when comparing two lysates.

However, since our experiments involve groups that do not necessarily have comparable total protein levels, this assumption may not hold. Similar to immunoprecipitation experiments with extensive washing of unbound material, normalising by median peptide counts can overemphasise background signals. Indeed, when we applied median peptide normalisation to the ASAP-ID^IT^ data, most known lamin interactors appeared “depleted” or not enriched, especially when using a threshold (FDR < 0.05, S0 = 1.7) to exclude proteins enriched in the control (**Fig. 2B**).

To find a more appropriate way to normalise the abundance of biotinylated proteins, we followed a previously described approach (15) that exploits the equal enrichment of four endogenously biotinylated proteins across experiments – these are proteins that are not expected to be directly influenced by the APEX2 reaction, but which should otherwise be co-purified in proportion to the starting material (input). These proteins are Acetyl-CoA Carboxylase Alpha (ACACA), Propionyl-CoA Carboxylase subunit Alpha (PCCA), Pyruvate Carboxylase (PC) and Methylcrotonyl-CoA Carboxylase subunit 1 (MCCC1) (16). The biotinylation of these four carboxylases is mediated by biotinoyl-5’-AMP (generated from biotin and ATP) (16), which is distinct from the chemistry of APEX2-mediated biotinylation, which uses biotin-phenoxyl radicals (generated from biotin-phenol and H_2_O_2_).

In a traditional proximity labelling experiment (APEX2-lamin fusion versus APEX2 alone), three of these enzymes appeared to be enriched at approximately the same level (**Fig. S4**), which is concordant with this hypothesis. However, ACACA appeared to have a higher level of enrichment in the APEX2-lamin eluates, suggesting that ACACA biotinylation is not independent of APEX2-lamin localisation. None of these proteins are known interactors of lamin, but we decided to exclude ACACA for normalising the data.

For the ASAP-ID^IT^ data and the traditional lamin-APEX2 proximity labeling data, normalisation to PC, PCCA, and MCC1 proteins yields results that more accurately reflect the levels of biotinylated proteins in total cell lysates via Western Blot and more effectively identify known interactors with lamin (**Fig. 2B & Fig. S5A-B**). Interestingly, while ACACA appeared unsuitable for normalisation of the traditional lamin-APEX2 fusion data (**Fig. S5C**), it seemed to produce a similar result to normalisation using PC, PCCA, and MCC1 for ASAP-ID^IT^ (**Fig. S5D**), indicating that the confounding effects of ACACA are context-dependent. As PC, PCCA, and MCC1 provided similar results for normalisation, we focused on further analysis using just the PC protein normalisation for simplicity, as it was the most abundant protein apart from ACACA of the four (**Fig. S6**).

Following PC protein normalisation, ASAP-ID^IT^ identified 19.8% of known lamin interactors as significantly enriched (3, 17) (45 out of 227) on the methanol-fixed cells, and 60.4% (137 out of 227) of them for the paraformaldehyde-fixed cells (**Fig. 2B**). This compared to the traditional lamin-APEX2 fusion proximity labelling experiment, which identified 16.7 % of the known lamin interactors (**Fig. 2B**). There was a strong correlation in enrichment of known lamin interactors between labelling (*in cis* or *in trans*) and fixation methods (**Fig. 2C**), and each experiment yielded a significant GO enrichment for nuclear envelope proteins (**Table S2**), providing confidence in the validity of the methodology. The methanol fixation did appear to have lower overall correlation to the other samples, which suggests some proteins do not fix as evenly in methanol as with paraformaldehyde. We also noticed that the known lamin interactors have higher protein intensities in the lamin-APEX2 fusion sample than the ASAP-ID^IT^ (**Fig. 2C**). This may reflect the more similar nature of the fusion experiment to the original BioID approaches that gave rise to the list of lamin interactors used in this study.

To demonstrate the application of ASAP-ID^IT^, we focused on profilin 1 (PFN1) and its mutants, which are associated with amyotrophic lateral sclerosis (ALS) (18). PFN1, which is 15 kDa, binds to monomeric actin through its actin-binding domain and has a role in regulating actin dynamics essential for various cellular processes, including cell motility, division and maintenance of cell shape (18). Beyond its interaction with actin, PFN1 binds to polyproline-rich sequences in proteins through its poly-L-proline (PLP) binding domain (19). PFN1 has other roles in gene regulation (^20, 21^), DNA damage response regulation (^22^), nuclear export of actin (^21^) and membrane trafficking (^23^).

We considered PFN1 (15 kDa) highly suitable for ASAP-ID^IT^ because a large tag like APEX2 (27 kDa) or TurboID (38 kDa) would be much bigger than the target protein being studied and therefore carry a higher risk of steric interference. In fact, we found that GFP (27 kDa) tagging of PFN1, caused the wild-type PFN1 to form abnormal aggregates (**Fig. S7A&B**).

Prior studies have found that four (untagged) ALS-related PFN1 mutants, C71G, M114T, E117G and G118V(18), formed distinct patterns of apparent aggregation compared to wild-type PFN1 when transfected into mammalian cells (24). The addition of the SunTag did not appear to change the localisation of PFN1 in HEK293T or HeLa cells when we compared our data (**Fig. 3A**) to published data (24). In essence, wild-type PFN1 formed a predominantly cytosolic pattern with no puncta or aggregates. E117G formed a similar pattern to wild-type PFN1, but with a higher proportion of PFN1 residing in the nucleus compared to the cytoplasm (24) (**Fig. 3B**). C71G, M114T and G118V all formed bright puncta characteristic of protein aggregates. The proportion of PFN1 in the nucleus compared to the cytoplasm increased, progressively, in order for C71G, to M114T, followed by G118V (**Fig. 3B**). These data suggest that subtle localisation differences and physical properties of the aggregates might impact what proteins are colocalised with each mutant.

**Figure 3.**
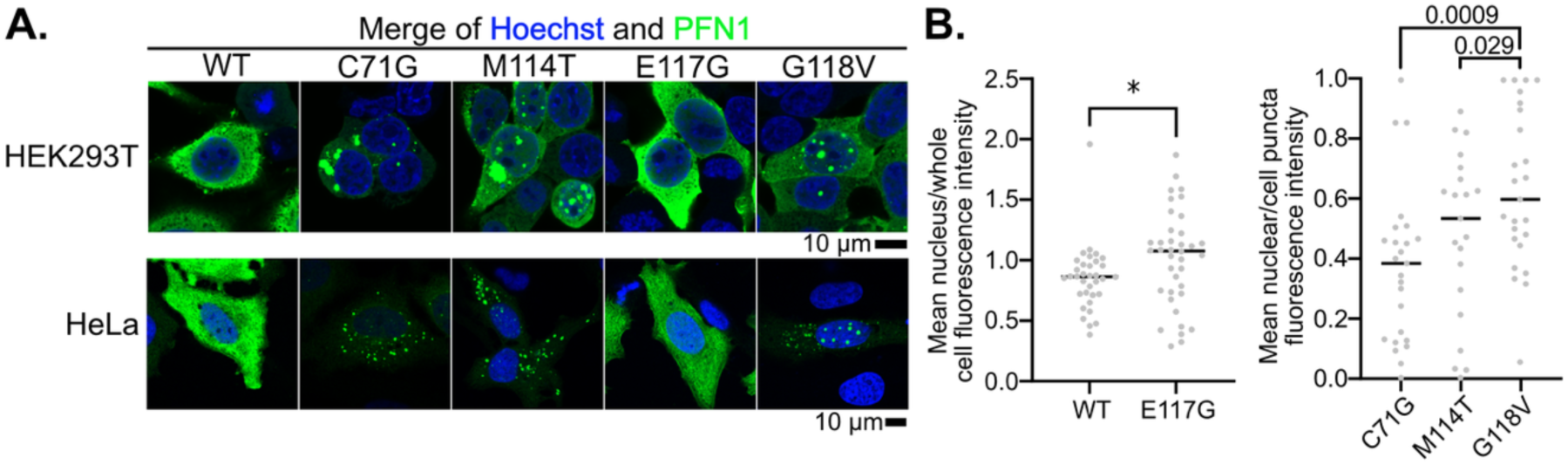
Localisation of PFN1 variants in cells. **A**. Immunostaining of the PFN1 variants fused to SunTag and HA in HEK293T cells and HeLa cells. The cells were then fixed with methanol and stained with anti-HA antibody. **B.** Quantification of PFN1 distribution in the HeLa cells. The mean intensity of WT and E117G PFN1 in the nucleus and whole cell was measured in the PFN1 variant-transfected HeLa cells by using ImageJ FIJI software. The ratio was calculated by dividing the mean PFN1 intensity in the nucleus by the mean intensity in the whole cell. T-test result shown: *, P<0.05. The number of puncta was measured in the C71G, M114T, and G118V PFN1-expressed HeLa cells using ImageJ FIJI software. The puncta ratio (nucleus/cell) was calculated by dividing the number of puncta measured in the nucleus divided by the number in the whole cell. Each dot represents one cell. Shown are the multiple comparison FDR q values of the Two-stage linear step-up procedure of Benjamini, Krieger and Yekutieli (11) from a one-way ANOVA (P=0.0034). The linear trend P-value was 0.008.

In total, 978 biotinylated proteins were enriched by proteomics upon application of ASAP-ID^IT^ to the PFN1 variants (**Table S3**; **Fig. 4A**). PFN1 was consistently more abundant when it had the SunTag, indicating that the proximity labelling had been effective. Of the proteins selectively enriched with WT PFN1 compared to the non-tagged PFN1, seven known interactors of PFN1 were found (EIF5A, IPO9, CDK12, STUB1, CTTN, FLNC, and CYFIP1). This result provided confidence that the proximity labelling had successfully detected the binding partners of PFN1. Five of these (EIF5A, IPO9, STUB1, CTTN, CYFIP1) were enriched with all PFN1 variants, indicating that the mutations do not disrupt their interaction with PFN1 (**Fig. S8A**). However, one of the ligand proteins, STUB1, was considerably more enriched in the C71G, M114T, and G118V variants suggesting a possible gain-of-function mechanism. Conversely, one protein, CTTN, showed a decrease in enrichment with the E117G, M114T, and C71G mutants, which may arise by a loss-of-function mechanism.

**Figure 4.**
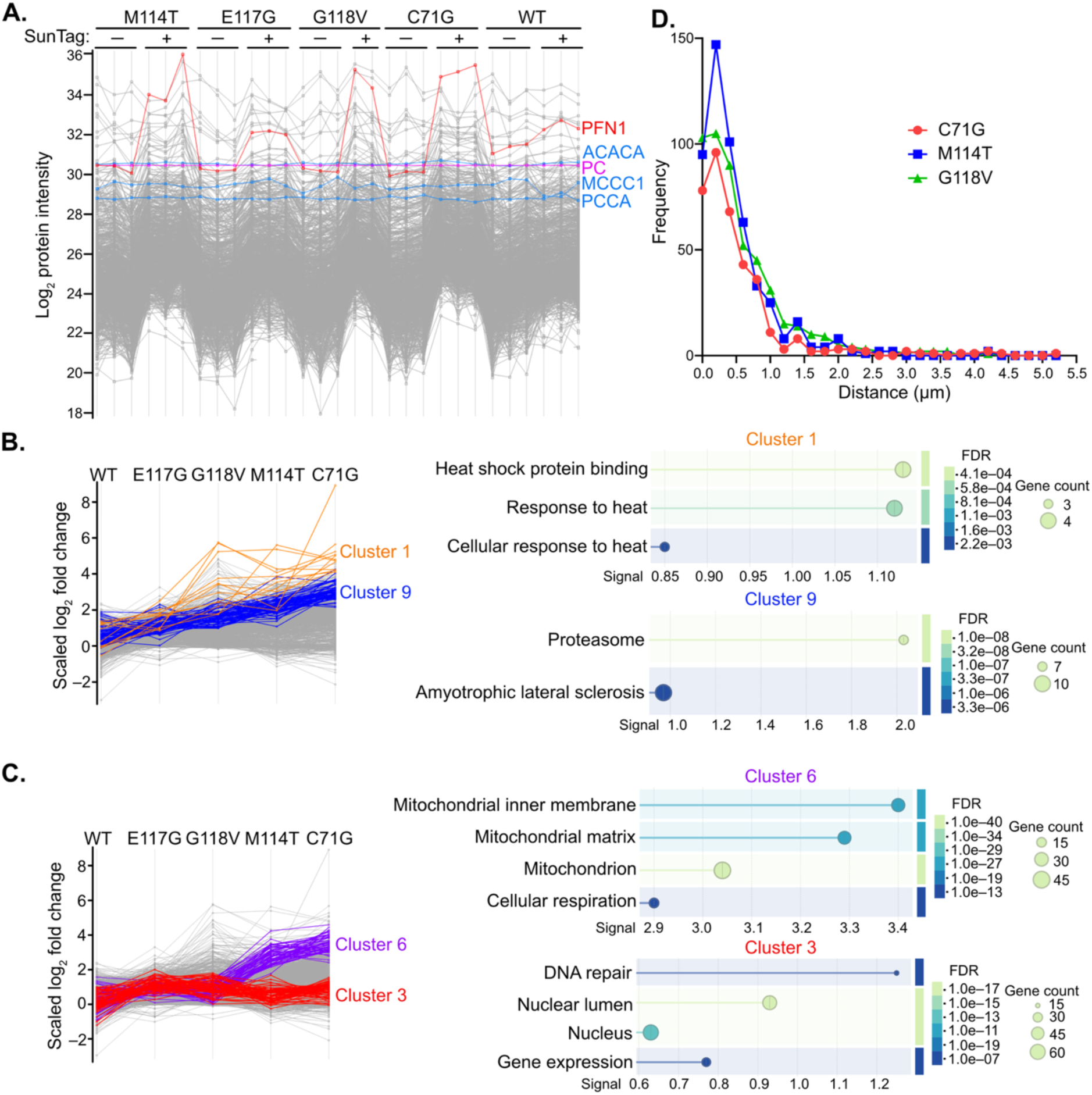
ASAP-ID^IT^ revealed interactomes of five PFN1 variants. **A**. Mass spectrometry intensity values of proteins detected across all SunTag-tagged and nontagged PFN1 variants. Data are shown after the filtering and imputation analysis and normalised to PC. The dots represent different proteins. **B**. Cluster 1 and cluster 9 derived from k-means clustering. Pathway enrichment analysis of cluster 1 and 9 was performed using STRING with default settings. The enrichment score (signal) represents a weighted harmonic mean of the observed-to-expected ratio and –log (FDR). Full details of enrichment patterns are provided in **Table S4**. **C**. Same logic as panel B, except showing cluster 3 and cluster 6. **D.** Histogram of shortest distances between PFN1 puncta in the cytosol and SDHA surfaces. Surfaces of PFN1 mutants and SDHA were reconstructed in 3D using Imaris, and shortest distances were measured between these surfaces within the software.

We postulated that proteins enriched with aggregated PFN1 would correlate with the mutants’ different localisation patterns. To examine this, we sorted the proteins into 13 clusters using k-means analysis (25) (**Fig. S8B** and **Tables S3 & S4**) and identified the patterns that emerged in the context of aggregation of the mutants based on imaging. Two of the clusters showed patterns consistent with proteins enriching with the aggregates (clusters 1 and 9) (**Fig. 4B**). Specifically, the enrichment of these proteins increased with increasing aggregation propensity of the PFN1 variants. Cluster 1 contained 12 proteins, including PFN1, FAF1, STUB1, DNAJC7, DNAJA1, NLN, POR, RBBP7, LARS2, CLPB, TOMM70 and IPO8. Four of these are heat shock binding proteins (DNAJA1, DNAJC7, STUB1 and FAF1) in GOMF: 0031072, enriched with an FDR of 0.0042 (**Fig. 4B)**. Cluster 9 included 66 proteins, which like cluster 1, had functions linked to protein quality control, specifically the proteasome (PSMC1, PSMC2, PSMC4, PSMC6, PSMD2, PSMD7, PSMD11) (in KEGG pathway hsa03050; enriched with an FDR of 1E-7) (**Fig. 4B)**. Collectively, these data suggest that aggregates recruit protein quality control machinery, which is consistent with what is known about protein aggregation more generally (26). Moreover, another well-known ALS-associated protein, TDP-43, was also in cluster 9, aligning with previous findings that mutant PFN1 can sequester endogenous TDP-43 (24).

Two other clusters stood out from the others. Cluster 3, with 69 proteins, showed selective enrichments for E117G and G118V mutants (**Fig. 4C**). In this cluster, 87% of these proteins are nuclear (GOCC:0005634, with enrichment FDR of 7.58E-14) (**Fig. 4C**). Wild-type PFN1 has functions in the nucleus, including roles in gene expression regulation (21), DNA replication (20), and DNA damage response and repair (22). In Cluster 3, 15 proteins involved in DNA repair (GOBP:0006281, with enrichment FDR of 1.87E-7), and 27 proteins involved in gene expression (GOBP:0010467, with enrichment FDR of 5.64E-7) were identified. These findings suggest that the E117G and G118V mutants may interfere with the DNA repair and gene expression machinery and may do this through an aggregation-independent mechanism, given that E117G doesn’t appear to aggregate.

Cluster 6 showed an enrichment of proteins for two mutants (M114T and C71G) over the others (**Fig. 4C**). It is noteworthy that these two mutants had a greater aggregation in the cytosol than the other PFN1 variants. 75% of the proteinsin cluster 6 (44 of 55) are mitochondrial proteins (GOCC:0005739 with enrichment FDR of 1.46E-33) (**Fig. 4C**). The preferential cytoplasmic localisation of these variants may explain this co-enrichment pattern. Few studies have previously linked PFN1 to mitochondria. One reported that wildtype PFN1 regulates mitochondrial morphology, dynamics, and respiration and that M114T PFN1 forms aggregates inside the mitochondria (27). Another reported a reduced mitochondrial content in M114T and C71G PFN1-transfected NSC-34 and HEK293T cells, but not in E117G and G118V PFN1-transfected cells(28). These findings agree with our data, suggesting that the M114T and C71G mutants localise or aggregate close to mitochondria where they influence various mitochondrial processes.

To investigate the localisation of PFN1, we conducted super-resolution fluorescence microscopy in HeLa cells and stained one of the mitochondrial proteins that we observed to be enriched with M114T and C71G, SDHA, which is located in the inner matrix compartment of the organelle where it is peripherally attached to the inner membrane, and co-stained the cells with PFN1 immunoreactivity. The surfaces of PFN1 aggregates and SDHA protein were reconstructed from z-stack images (**Fig. S9**). The shortest distance between each PFN1 aggregate and SDHA was measured to evaluate their co-localisation. Strikingly, most of the cytosolic puncta of M114T and C71G are not within the mitochondria but are proximal to it; more than half of the puncta were located within 0.5 µm of SDHA (**Fig. 4D**). Notably, SDHA was in cluster 6, it is more enriched in the M114T and C71G mutants than the others. These results suggest that ASAP-ID labelling is not only capable of detecting changes in the proximity of different PFN1 mutants but also that the labelling radius may extend to the micrometre scale.

## Discussion

Here, we developed a new proximity labelling method, ASAP-ID, that removes the requirement to fuse APEX2 or TurboID to the protein of interest genetically. This offers a great benefit in reducing the steric impact on the function of the protein of interest. ASAP-ID has some similarities to other methods that couple biotinylation enzymes to antibodies targeting the protein of interest or the post-translational modification (29, 30). The benefit of such approaches over ASAP-ID is that endogenous proteins can be directly targeted, which enables samples from patients or other organisms to be assayed. However, the approaches depend on the quality of the antibody and its specificity, and it may be difficult to control for background. In principle, ASAP-ID^IT^ could be developed to work on untagged endogenous proteins by fusing nanobodies or scFvs to the APEX2. Another use case is on tissue from mouse models harbouring whole body knock-in of a SunTag to a protein of interest. A similar approach using the similarly sized FLAG tag and classical immunoprecipitation to study tissue-specific differences in protein composition of the mitochondrial ribosome (31).

One of the questions that arose from this work is how far the proximity labelling reaches beyond the target protein in ASAP-ID. This comes from the large number of interactors detected for lamin (>300) and the fact that for PFN1 there was substantial labelling of mitochondrial proteins when it was clear by microscopy they were in physically distinct locations. The labelling radius for APEX2 reaction products is determined by both the half-life of the radical and the concentration of the quencher, glutathione, in the environment (32). The predicted labelling radius of the APEX2 under cellular settings is 20 nm (4). 20 nm is lower than the resolution limit of the super-resolution imaging in our experiments. Hence, it is more likely that the limit is much larger than this, possibly more than 100 nm. The diffusion of free radicals has been observed for the HRP-based TSA-seq method (33), which suggested a limit of 1 µm, which is more consistent with our observations. Radii of up to 0.55 µm have also been reported in paraformaldehyde-fixed cells expressing APEX2 (34).

Another contributing factor to the distance of labelling may be the size of the protein complex in the ASAP-ID (epitope bound to antibody-APEX2 fusion). While the epitope is small and directs the scFv to bind to the first 10 amino acids of the SunTag (EELLSKNYHL) (12), the scFv creates a “linker” to the fused APEX2 that extends the labelling radius. The proximity labelling range has been previously reported to increase through the addition of a 25-nm linker consisting of 13 repeats of GGGGS in genetic fusions between BirA variant BioID2 and a target protein, Nup43 (35). In this example, the extended linker allowed a greater number of proteins within the same complex to be captured, but also allowed a far greater number of other proteins to be detected, which is consistent with our observations with lamin and profilin (35). The scFv has a molecular weight of approximately 26.5 kDa, which corresponds to approximately 3 nm in width (36), which is far shorter than the disordered 13×GGGGS repeat. A further factor that could influence labelling radii is the residual level of glutathione. In cells, glutathione is present in millimolar concentrations and acts as an antioxidant (37). It can reduce free radicals, such as hydroxyl ions, and therefore quench the APEX2 biotinylation. However, upon fixation, most of the glutathione is expected to be oxidised (38) or washed away (39), which may attenuate the quenching effect, and in turn permit a more spatially promiscuous APEX2 reaction.

For applications of ASAP-ID there are several important considerations to note. For the *in-cis* co-expression approach of ASAP-ID^IC^, the constant expression and binding of the nanobody-APEX2 to the epitope would be expected to confer steric interference, not dissimilar to the traditional proximity methods involving the fusion of APEX2. However, it would offer the benefit of capturing newly synthesised proteins more effectively than direct fusion because the binding of the nanobody-APEX2 to the epitope would occur independently of the requirement of APEX2 to fold. ASAP-ID^IT^ overcomes these steric effects, but the extra steps for fixation and protein harvest may lead to a loss of protein identifications. For example, methanol may not fix all proteins evenly and hence there could be leaching of some proteins, notably hydrophobic proteins (40). Paraformaldehyde cross-linked molecules require heat-based retrieval steps to reverse the cross-links. Unevenness in the efficiency of different cross-linked moieties may lead to biases in protein abundances that are recovered. Overall, our data showed that paraformaldehyde-fixation provides data that more closely correlates with the traditional proximity approach (APEX2-lamin) than methanol-fixation, which suggests that paraformaldehyde better preserves the representation of proximal proteins.

Our application of ASAP-ID^IT^ on PFN1 demonstrated the power of the approach, enabling us to highlight how mutations linked to ALS can lead to differences in endogenous protein coaggregation. When considering the mutations on their pure tendency to aggregate, ASAP-ID^IT^ showed a correlation between aggregation propensity and enrichment with protein quality control machinery, including DNAJA1, DNAJC7, STUB1, FAF1, the PSMC family, and the PSMD family. These proteins are also linked to protein aggregation in other studies (41–44). Both DNAJA1 and DNAJC7 are members of the J-domain protein (JDP) family, which functions as co-chaperones with Hsp70 proteins, aiding in protein folding and preventing aggregation (45). It has been shown that DNAJC7 preferentially binds and stabilises natively folded tau protein, which is a pathological protein in Alzheimer’s disease, and prevents tau conversion into amyloids (41). DNAJA1 has been shown to have a similar but lower effect on tau aggregation than DNAJC7 (41). Moreover, the loss of DNAJC7 is a genetic risk factor for ALS (46). STUB1 encodes the C-terminus of Hsc70-Interacting Protein (CHIP), which functions as both a co-chaperone and an E3 ubiquitin ligase. CHIP interacts with chaperones such as Hsp70 and Hsp90, facilitating the ubiquitination and subsequent proteasomal degradation of misfolded proteins, thereby maintaining protein homeostasis (47). STUB1 exhibits neuroprotective effects by regulating the degradation of mutant SOD1 (42). Additionally, it is reported to interact with PFN1, ubiquitinating it for proteasome-mediated degradation (43). Hence, its accumulation with aggregated PFN1 may reflect a protective role for STUB1 in the clearance of misfolded protein aggregates. FAF1, also known as Fas-associated factor 1, is a ubiquitin receptor that contains multiple ubiquitin-related domains. The accumulation of FAF1 has been observed to play a key role in dopaminergic neuronal degeneration, suggesting its involvement in the pathogenesis of Parkinson’s disease through mechanisms related to protein aggregation (44). The PSMC family comprises ATPase subunits in the 19S regulatory particle of the 26S proteasome, whereas the PSMD family contains non-ATPase subunits in the 19S regulatory particle of the 26S proteasome (48). Both families play important roles in the proteasome’s function of degrading misfolded or damaged proteins (48, 49). Impairment of the ubiquitin-proteasome system has been demonstrated to result in the accumulation of mutant PFN1 aggregates (18). Cells expressing PFN1 mutants C71G, M114T, and G118V exhibited numerous large protein inclusions following treatment with the proteasome inhibitor MG132 (18). Cells expressing the E117G mutant displayed a moderate level of aggregation, while those expressing wild-type PFN1 showed minimal aggregate formation (18). These findings, in conjunction with our data, suggest that PFN1 aggregates may sequester components of the protein quality control machinery, thereby compromising proteostasis and contributing to the persistence and accumulation of misfolded protein aggregates.

The other more intriguing result was the pattern of aggregate location in the cell. The C71G and M114T PFN1 mutants were more likely to be enriched with mitochondrial proteins, whereas the E117G and G118V mutants were more enriched with nuclear proteins. This can be explained by the preferred localisation of PFN1 aggregates in cells; the C71G and M114T have a higher proportion of aggregates in the cytoplasm, whereas more aggregates of the G118V mutant appear in the nucleus.

The selectivity of aggregate location may play a role disease pathomechanisms. The potential impact of this is well illustrated by the aggregation behaviour of ALS-associated protein Fused in Sarcoma (FUS) (50). FUS normally resides in the nucleus where it operates in functions related to transcription, mRNA splicing, and transport (51). In its role regulating transcription, FUS binds DNA, recruits RNA polymerase II, and modulates transcriptional activators (52). In its role in regulating mRNA splicing, FUS binds to heterogeneous nuclear ribonucleoproteins (hnRNPs) and the U1 small nuclear ribonucleoprotein (snRNP) complex (53, 54). In ALS patient-derived fibroblasts, FUS can form nuclear aggregates, which are associated with disruptions in RNA polymerase II function (55). Mutations in the C-terminal region of FUS that cause ALS, can instead, mediate mislocalisation and aggregation in the cytoplasm (56). In this context, the cytoplasmic FUS aggregates sequester RNA-binding proteins, leading to impaired RNA granule formation (57) and aberrant RNA splicing (56). Collectively, these findings illustrate the potential for nuclear or cytoplasmic aggregation to drive different mechanisms of gain of toxic functions.

Our data with PFN1 suggests that differential localisation of aggregation may selectively modulate PFN1 dysfunction in DNA replication and DNA repair (20, 22), or mechanisms involving mitochondrial function (27).

Also of relevance to ALS mechanisms was the identification of TDP-43 co-aggregating with PFN1 mutants in a manner correlated with PFN1 aggregation propensity (TDP-43 was in cluster 9). TDP-43 cytoplasmic mislocalisation from a normal nuclear localisation is a hallmark of ALS pathology, and the data here, as well as elsewhere, suggest PFN1 can influence this behaviour (24).

In summary, ASAP-ID offers a flexible format that allows for the use of various biotinylation enzymes (TurboID and APEX2), epitope tags (SunTag and MoonTag), and labelling methods (methanol fixation or paraformaldehyde fixation for *in trans* labelling, as well as the in cis labelling). All of these approaches have been demonstrated here to label the proximity protein of the model protein human lamin A in the immunostaining assay. These properties will make ASAP-ID, particularly the adaptability and flexibility of ASAP-ID^IT^, a standout approach for versatile proximity labelling in cells and tissue samples.

## Acknowledgements

Imaging was conducted with support from the Biological Optical Microscopy Platform (BOMP) and the Materials Characterisation and Fabrication Platform (MCFP) at the University of Melbourne. The proteomics data was collected by the Mass Spectrometry and Proteomics Facility at Bio21 Institute, The University of Melbourne. The work was funded through grants to DMH (Australian Research Council: DP250100240 and DP230101050). DAS is supported by a National Health and Medical Research Council Investigator Fellowship (GNT2009732). This work was supported in part by grants from the US Air Force Office of Scientific Research (FA9550-20-1-0241) and the St. Jude Research Collaborative on the Biology and Biophysics of RNP granules to RVP.

Data are available via ProteomeXchange with identifier PXD066625.

This article contains supplemental data and figures

## Notes

### Competing Interest Statement

The authors have declared no competing interest.

