## Supplemental Figures for "ASAP-ID: Proximity labelling with small tags"

### Supplementary figures

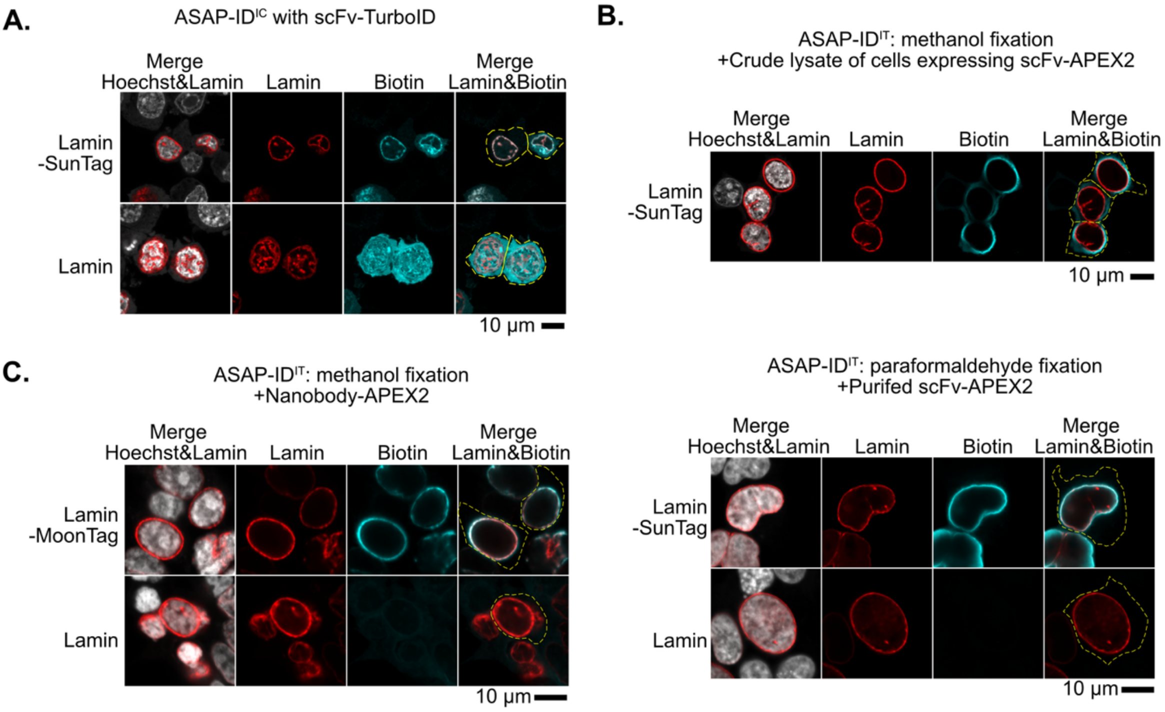


**Supplementary Figure 1. ASAP-ID can label lamin proximal proteins with different approaches, biotinylation enzymes and epitope tag. A.** ASAP-ID$^{IC}$ with TurboID and SunTag. The constructs shown on the left were co-transfected with scFv fused to TurboID in HEK293T cells. Cells were then subjected to biotinylation and fixed with methanol. **B.** ASAP-ID$^{IT}$ approaches. HEK293T cells transfected with lamin constructs (shown on left) and then fixed with methanol or paraformaldehyde, and then treated with the scFv-APEX2 fusion proteins as shown. Antibody-APEX2 was either purified using HA antibody affinity chromatography or expressed in cells, where cell lysates were added. **C.** MoonTag mediated ASAP-ID$^{IT}$. HEK293T cells expressed lamin constructs (shown on left) and were then fixed with methanol. Purified nanobody-APEX2 was added to fixed cells.

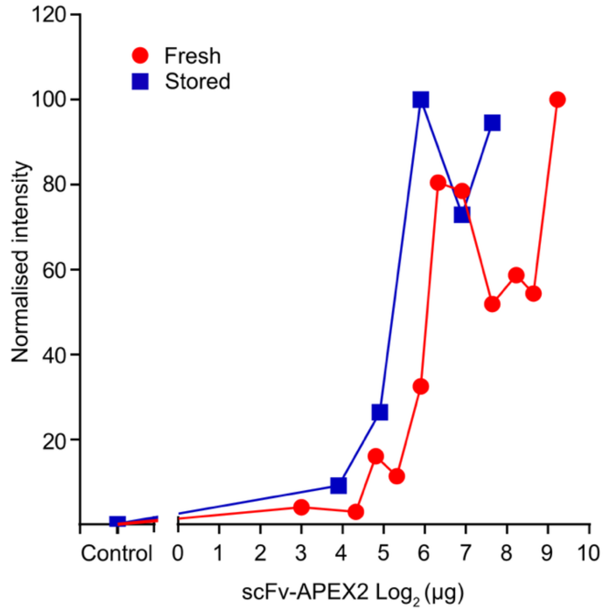


**Supplementary Figure 2. Activity of purified APEX2 antibody. A.** Shown are fluorescence values of biotin stain after ASAP-ID$^{IT}$ on HEK293T cells transfected with SunTag-lamin-mCherry, and then treated with purified scFv-APEX2. Each value was derived from the average fluorescence of 12 cells. The normalised intensity was calculated by dividing the biotin immunostain fluorescence intensity by the mCherry intensity. The scFv-APEX2 was produced by immunoaffinity capture from transfected HEK293 cells. Freshly purified and snap frozen, thawed after one month storage at –80 °C, are shown.

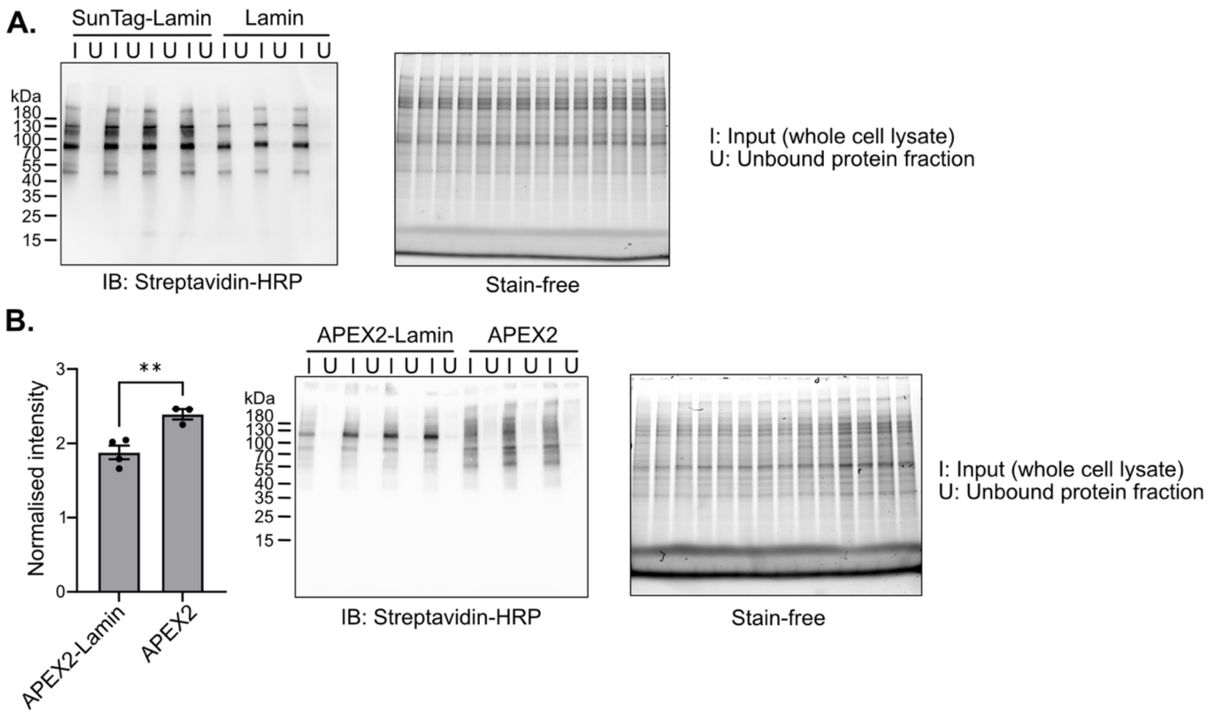


**Supplementary Figure 3. Biotinylation levels in the ASAP-ID and traditional proximity labelling approach (Lamin-APEX fusion). A.** Western Blot analysis of HEK293T cells expressing lamin and treated by ASAP-ID$^{IT}$. The biotinylated proteins were detected by the streptavidin-HRP. The unbound protein fraction was the result of a streptavidin bead pull-down of the input. **B.** Same logic as panel A, except the samples reflect APEX2-lamin fusions instead of the ASAP-ID$^{IT}$ protocol. In this case the negative control is APEX2 alone expressed in the HEK293T cells. Graphs show means ± SEM of the digitised Western Blot data. T-test result shown: **, P-value = 0.009.

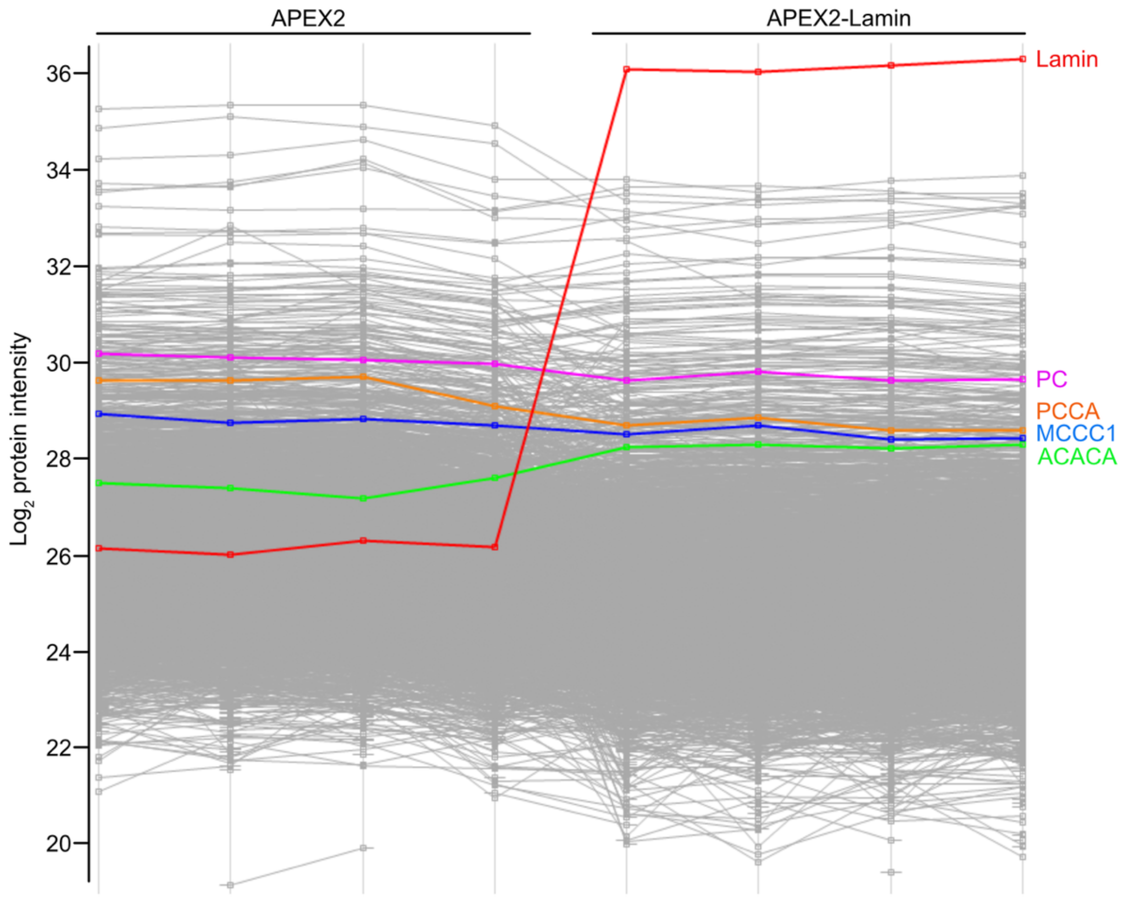


**Supplementary Figure 4. Abundance profile of proteins identified in the APEX2-lamin proteomics.** In the experiment, proximity labelling was undertaken comparing HEK293T cells expressing lamin fused to APEX with APEX alone. Data points show individual protein abundances after the filtering and imputation analysis of four replicates.

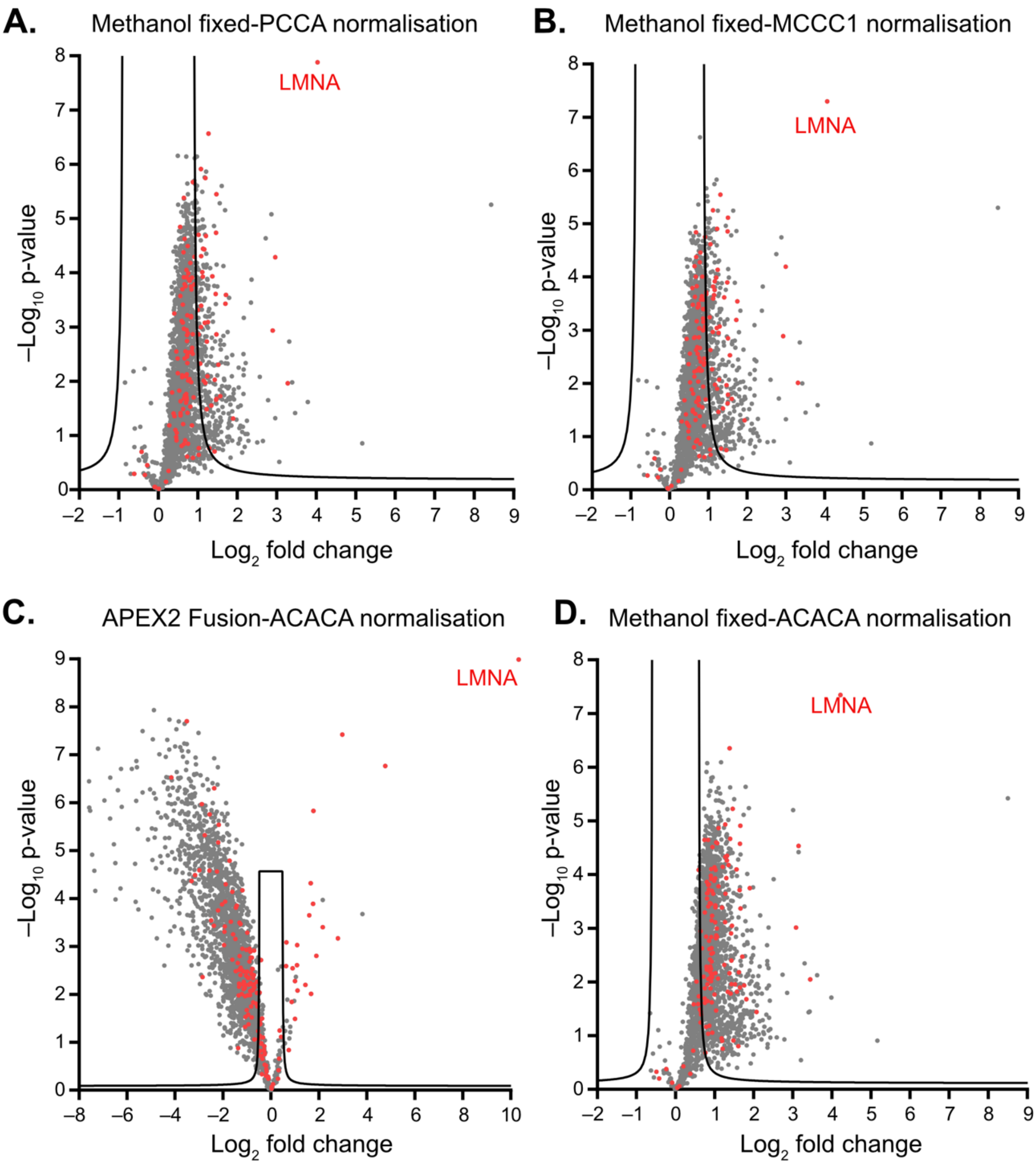


**Supplementary Figure 5. Comparison of different normalisation approaches to analyse the ASAP-ID and direct fusion methods. A**. ASAP-ID$^{IT}$ on cells transfected with lamin. Data was normalised to protein PCCA abundance. Data shows lamin-SunTag/lamin changes. Red dots represent previously established lamin interactors. Threshold was set as FDR < 0.05 and S0 = 2. **B**. Same logic as panel A, except the normalisation was based on protein MCCC1. **C**. Traditional proximity labelling experiment data comparing Lamin-APEX fusions with APEX alone with data normalised to ACACA protein levels, **D.** Same logic as panel A, except the normalisation was based on protein ACACA.

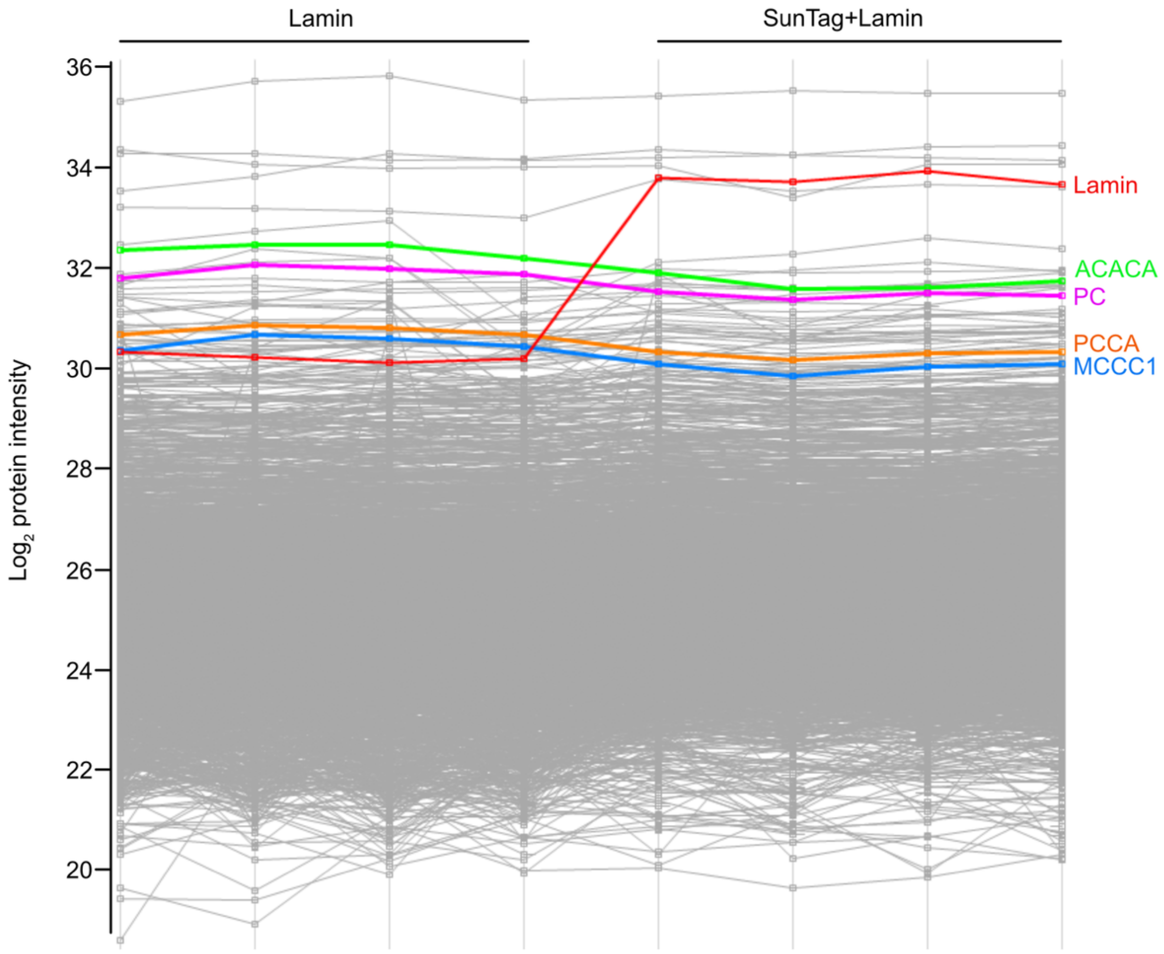


**Supplementary Figure 6. Protein abundance profiles of proteins identified in ASAP-ID$^{IT}$ using methanol fixation.** HEK293T cells were transfected with lamin-SunTag fusion or untagged lamin. Data points show individual protein abundances after the filtering and imputation analysis of four replicates.

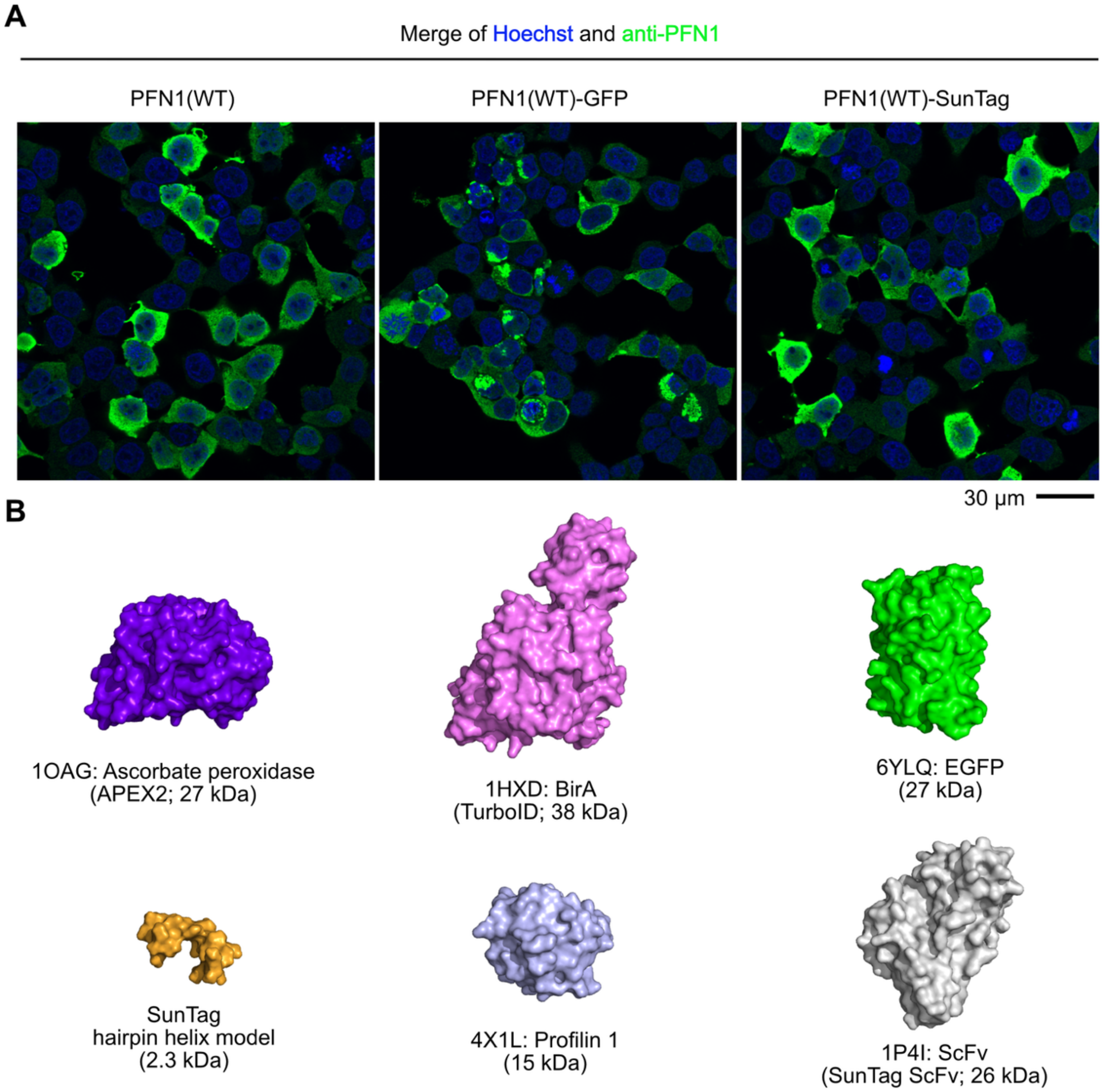


**Supplementary Figure 7. Comparison of the expression of different tagged wild-type PFN1. A.** Immunostaining of the GFP-tagged, SunTag-tagged tagged and nontagged wild-type PFN1. HEK293T cells were transfected with different constructs of tagged PFN1, cells were then fixed with paraformaldehyde and stained with anti-PFN1 antibody. **B.** Renders of proteins for consideration of scale. Protein models are shown with relevant PDB entries, and are shown to scale. Rendering was performed by PyMol (58).

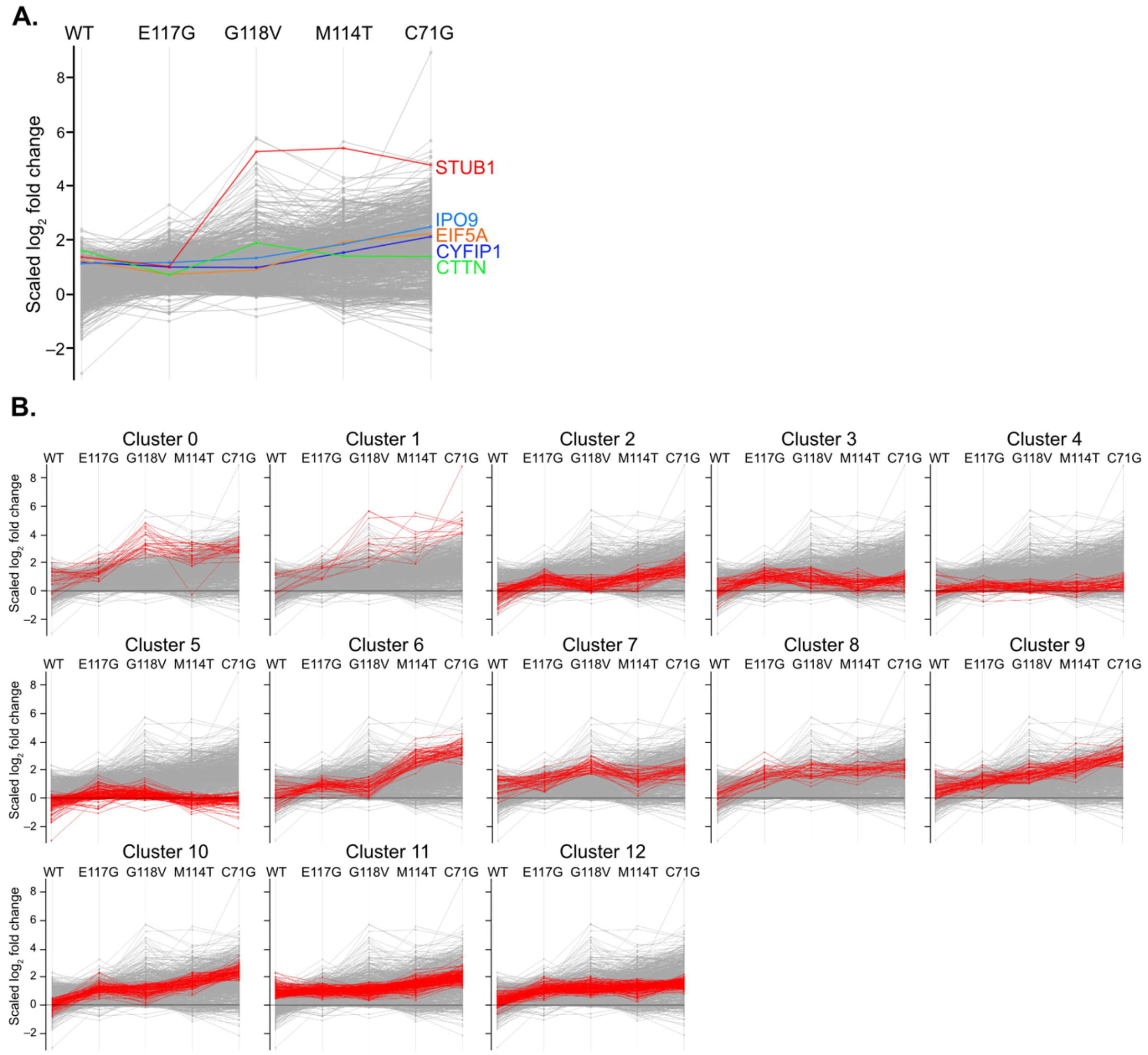


**Supplementary Figure 8. K-Means cluster profiles of proteins identified in the PFN1 ASAP-ID[IT] proteomics. A.** The abundance profiles of selected known PFN1-interacting proteins. Data points indicate different protein abundances after smoothing of replicates using PERCEPT to highlight the important changes (refer to methods). **B.** The complete list of 12 clusters (in red) generated by the k-means clustering analysis.

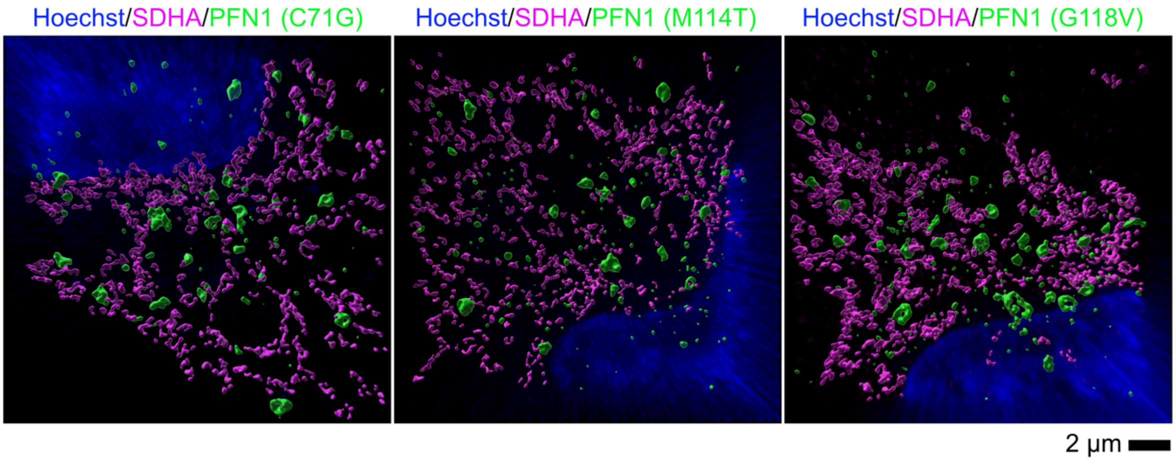


**Supplementary Figure 9. Super-resolution microscope images of PFN1 aggregates and SDHA.** HeLa cells were transfected with HA-tagged PFN1 mutants. Cells were fixed with methanol and stained with anti-SDHA and anti-HA antibodies to detect the SDHA and PFN1 in cells. Super-resolution images were acquired with a Zeiss Elyra 7 Lattice SIM super-resolution microscope, and the surface of the protein was built using Imaris.
